# Beneficial Effects of Celastrol on Immune Balance by Modulating Gut Microbiota in Dextran Sodium Sulfate-Induced Ulcerative Colitis

**DOI:** 10.1101/2021.09.28.462065

**Authors:** Mingyue Li, Weina Guo, Yalan Dong, Wenzhu Wang, Chunxia Tian, Zili Zhang, Ting Yu, Haifeng Zhou, Yang Gui, Kaming Xue, Junyi Li, Feng Jiang, Alexey Sarapultsev, Shanshan Luo, Heng Fan, Desheng Hu

## Abstract

Ulcerative colitis (UC) is a chronic inflammatory bowel disease caused by multi-factors including colonic inflammation and microbiota dysbiosis. Previous studies have indicated that Celastrol (CSR) has strong anti-inflammatory and immune-inhibitory effects. Here, we investigated the effects of CSR on colonic inflammation and the mucosal immunity in an experimental colitis model, and addressed the mechanism by which CSR preforms the protective effect. We characterized the therapeutic effects and the potential mechanism of CSR in treating UC using histological staining, intestinal permeability assay, cytokine assay, flow cytometry, fecal microbiota transplantation (FMT), 16S rRNA sequencing, untargeted metabolomics, and cell differentiation approaches. CSR administration significantly ameliorated DSS-induced colitis, as evidenced by the recovery of body weight and colon length, decreased disease activity index (DAI) score, as well as decreased intestinal permeability. CSR down-regulated the secretion of proinflammatory cytokines, upregulated the anti-inflammatory mediators, and improved the balances of Treg/Th1 and Treg/Th17 to maintain colonic immune homeostasis. However, the protective effects of CSR disappeared when the antibiotic cocktail was applied to deplete the gut microbiota, and the gut microbiota-mediated effect was confirmed by FMT. Furthermore, CSR treatment increased the gut microbiota diversity and composition, and raised the metabolic productions of pyruvate and adenosine, which probably involve in gut microbiota mediated protective effect. In conclusion, CSR ameliorates colonic inflammation in a gut microbiota-dependent manner. The underlying protective mechanism is associated with the rectified Treg/Th1 and Treg/Th17 balance, and increased pyruvate and adenosine production. The study provided the solid evidence that CSR might be a promising therapeutic drug for UC.

## Introduction

Ulcerative colitis (UC) is one of two major forms of inflammatory bowel disease and involves diffuse inflammation of the colonic mucosa. With the poor quality of life, high morbidity and risk of colitis-related colorectal cancer, UC has become a public health threat[1, 2]. The current treatment strategy for UC is based on the severity, distribution, and pattern of disease, including the use of oral and/or topical 5-aminosalicylic acid as the first-line medication for induction and maintenance therapy, nonsystemic and systemic corticosteroids, or a combination of anti-tumor necrosis factor and immunomodulator therapy[3, 4]. Due to the low remission rate of new drugs or the secondary loss of response, the patients unresponsive to these therapies suffer from high morbidity associated with delayed surgery[5].

The pathogenesis of UC still remains unclear. Several factors, including the genetic susceptibility, dysregulated immune responses, imbalance of gut microbiota, and environmental exposure have been reported to be implicated[6]. The imbalance between T helper (Th) cells and regulatory T (Treg) cells has been proved to be correlated with the occurrence and severity of UC[7]. During UC progression, Th1 and Th17 cells are usually increased and exert pathogenic effects by releasing the proinflammatory cytokines including IFN-γ and IL-17A, respectively, while Treg cells, which inhibit Th1 and Th17 activities through intercellular communication, are decreased[8–13].

Dysbiosis of gut microbiota could skew the balance of Treg/Th17 cells, and then trigger exaggerated inflammatory responses in UC[14]. It has been reported that the composition of the gut microbiota in patients with UC markedly differs from those of healthy individuals[15]. Furthermore, the dysbiosis between probiotics and pathogenic bacteria, and the reduction of bacterial diversity play an important role in the development of UC[16, 17]. In addition, the gut microbiota-derived metabolites function as an intermediate link between microbiota and local immune responses. For instance, it has been confirmed that accumulated short-chain fatty acid can alleviate colitis by regulating Treg/Th17 balance[18–20]. Therefore, maintaining the homeostasis of gut microbiota might be a promising therapeutic strategy for UC.

Celastrol (CSR), isolated from the root xylem of *tripterygium wilfordii* (TW), a Chinese herb used in Traditional Chinese Medicine for several centuries. With strong antioxidant, anti-inflammatory, and anticancer properties, an increasing number of studies were published highlighting the medicinal usefulness of celastrol in diverse clinical areas, namely, rheumatoid arthritis, systemic lupus erythematosus, inflammatory bowel diseases, osteoarthritis and allergy, as well as in cancer, neurodegenerative disorders[21]. In mice models of DSS-induced colitis, it has been found that celastrol ameliorates acute intestinal injury and prevents the loss of intestinal epithelial homeostasis through the reduction of colonic oxidative stress, inhibition of NLRP3-inflammasome and IL-23/IL-17 pathway, reduction of inflammatory cytokines and increase in IL-10 and TNF levels, attenuation of neutrophil infiltration and upregulation of E-cadherin expression[22, 23].

However, the intrinsic therapeutic mechanism of CSR on UC still needs to be clarified. Given the vital role of gut microbiota dysbiosis and immune dysregulation in the pathogenesis of UC, we sought to address the potential impacts of CSR on gut microecosystem and immune system in dextran sodium sulfate (DSS)-induced colitis mouse model.

## Results

### Celastrol administration ameliorated DSS-induced colitis

To investigate the therapeutic effect of CSR (**Figure S1A**) on UC, DSS-induced colitis model was established in this study. Mice were treated with 3.0% DSS in the drinking water, and CSR suspended in saline was administered by oral gavage for 7 days (**Figure 1A**). Compared with the DSS group, CSR administration significantly ameliorated DSS-induced colitis, as evidenced by the markedly reduced body weight loss (**Figure 1B**), and improved colon length (**Figure 1C**). Furthermore, DAI score based on the assessment of stool consistency, bloody stool, and weight loss concorded with the above results, validating the curative effect of CSR (**Figure 1D**). To test the intestinal permeability, the mice were gavaged with FITC-dextran 4 hours before being euthanized. Compared to control group, mice with TG treatment exhibited decreased serum level of FITC-dextran, indicating the relatively intact epithelial barrier (**Figure 1E**). Additionally, H&E staining were performed to evaluate the extent of injuries of colon. Compared with DSS-treated mice, the distorted crypts, loss of goblet cells, epithelial injury, and the infiltration of inflammatory cells in the mucosa and submucosa were dramatically alleviated in mice of the DSS+CSR+ group. These observations indicated that CSR well controlled colon inflammation and maintained epithelial barrier integrity (**Figure 1F**), which was further confirmed by Alcian blue staining showing the intensity of goblet cells (**Figure S1B**). Since tight junction plays a vital role in maintaining the barrier of epithelium in colon, next we assessed the expression of tight junction related proteins. Data revealed that DSS inhibited the expression of *Occludin*, *Cdh1*, *Zo-1* and *Muc2*, which were rescued by CSR treatment (**Figure 1G**).

**Figure 1.**
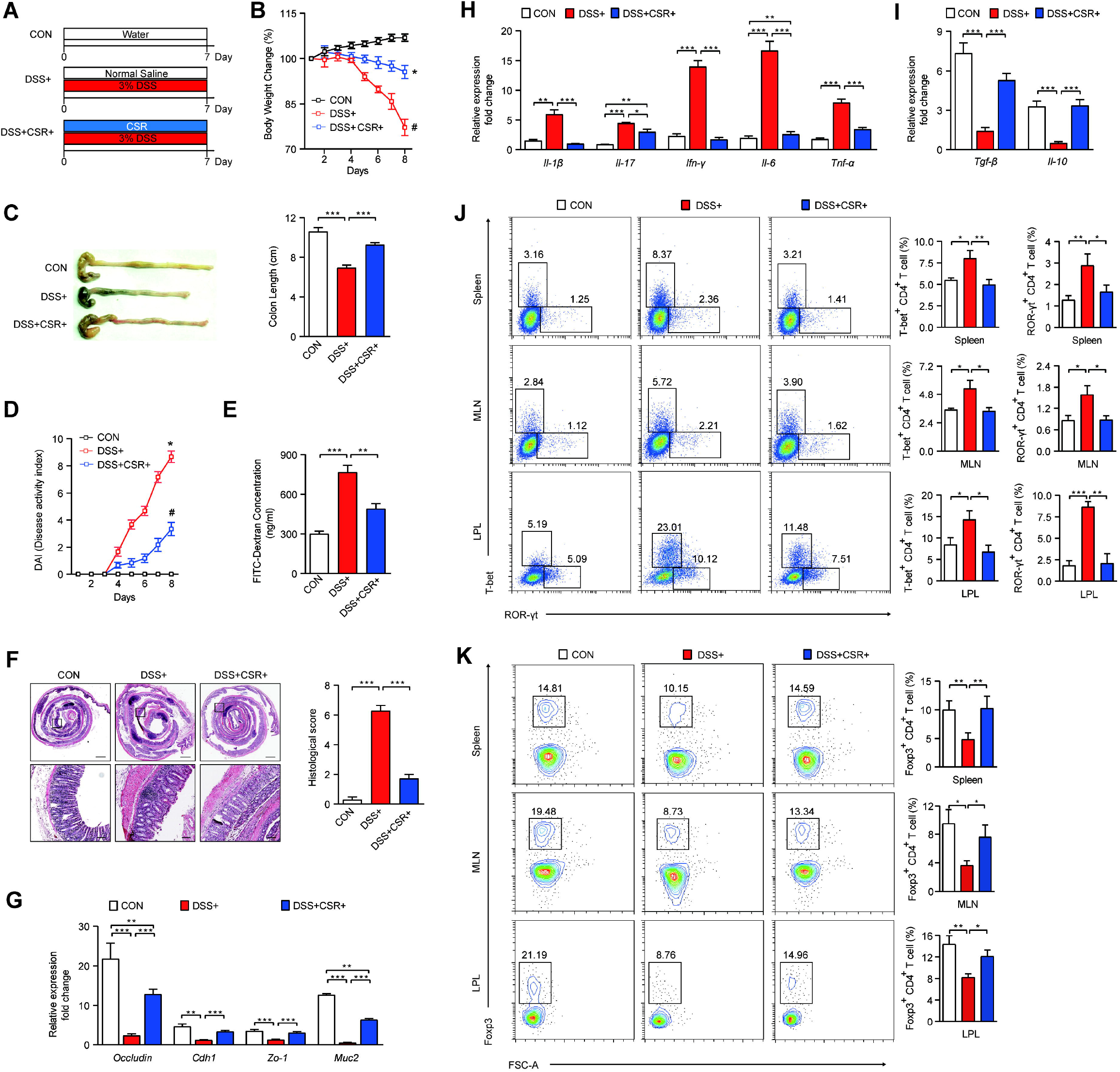
CSR attenuated DSS-induced experimental colitis in mice. **(A)** Schematic diagram illustrates the experimental design. (B) Body weight percentage changes of each group (n=10). (C) Measurement of the length of colons harvested from mice in each group (n=10). (D) The effect of CSR on DAI in mice (n=10). (E) Mice in each group received an oral gavage of FITC-dextran (0.5mg/g) and serum FITC-dextran concentrations were determined 4□h later (n=5). (F) H&E staining (Bar=1mm above, Bar=100um below) sections and histological scores of colon tissue from mice in each group (n=5). (G) The expression levels of *Occludin*, *Cdh1*, *Zo-1* and *Muc2* of colon tissue in each group (n=5). (H) The expression levels of pro-inflammatory cytokines of colon tissue in each group (n=5). (I) The expression levels of anti-inflammatory cytokines of colon tissue in each group (n=5). (J) T-bet^+^CD4^+^ (Th1) cells and ROR-γt^+^CD4^+^ (Th17) cells in spleen, MLN and LPL from the control, DSS+ group and DSS+CSR+ group were analyzed by flow cytometry and bar charts of the percentage of Th1 and Th17 cells (n=5). (K) Foxp3^+^CD4^+^ (Treg) cells in spleen, MLN and LPL from the control, DSS+ group and DSS+CSR+ group were analyzed by flow cytometry and bar charts of the percentage of Treg cells (n=5). Data are presented as mean ± *SEM*. \**P* < 0.05, significantly different as indicated.

Immune cells and secreted inflammatory cytokines are important mediators for immune homeostasis in the gut. Hence, we examined these mediators in the colon tissues. The proinflammatory cytokines, including *Il-1β*, *Il-17a*, *Ifn-γ*, *Il-6*, and *Tnf-α* were down-regulated, while the anti-inflammatory mediators *Tgf-β* and *Il-10* were up-regulated in colon tissues of mice in the DSS+CSR+ group (**Figure 1H and 1I**). The frequencies of IFN-γ^+^CD4^+^ and IL-17A^+^CD4^+^ T cells in spleen and MLN in DSS+CSR+ group were reduced (**Figure S1C**). Meanwhile, in spleen, MLN and colonic LPL, the percentages of T-bet^+^CD4^+^ and ROR-γt^+^CD4^+^ T cells were down-regulated, and Foxp3^+^CD4^+^ T cells were up-regulated in DSS+CSR+ group mice (**Figure 1J and 1K**). Together, these results indicated that CSR can prevent DSS-induced colitis.

### Celastrol up-regulated the differentiation of Treg cells in vitro

To further explore whether CSR directly regulate T cell differentiation, naive CD4^+^ T cells were isolated from spleen of WT mice, and were co-cultured with different concentration of CSR under the corresponding conditions for Th1, Th17 and Treg cells differentiation in vitro. The results showed that CSR had no effect on the differentiation of IFN-γ^+^CD4^+^ T cells and IL-17A^+^CD4^+^ T cells in the indicated CSR concentration. However, Foxp3^+^CD4^+^ cells were elevated when the concentration of CSR is 4 μmol/L (**Figure S2**).

### Celastrol alleviated colitis in a gut microbiota-dependent manner

To investigate whether gut microbiota mediated the protective effect of CSR on DSS-induced colitis, the WT mice were treated with quadruple antibiotic cocktail (ABX) for gut microbiota depletion before DSS treatment (**Figure 2A**). Following the 3-week ABX treatment, the gut microbiota *Eubacteria*, *Lactobacilius*, *MIB and Eubacterium rectale-Clostridium coccoides* were significantly decreased in fecal samples of ABX-treated mice (**Figure S3A**), which were consistent with previous research. Moreover, there was no difference in the morphological of liver, kidney, intestine, or colon, and the serum levels of ALT, AST, BUN, and CRE after ABX treatment compared to untreated mice, indicating the non-organ toxicity of ABX (**Figure S3B and S3C**). Impressively, the theraputic effect of CSR was reversed after gut microbiota depletion. ABX+DSS+ group mice and ABX+DSS+CSR+ group mice displayed no significant difference in the following index, such as body weight loss (**Figure 2B**), colon length (**Figure 2C**), DAI score (**Figure 2D**), intestinal permeability (**Figure 2E**), histological changes and scores (**Figure 2F**), goblet cells(**Figure S4A**), tight junction proteins (**Figure 2G**), proinflammatory cytokines and anti-inflammatory mediators (**Figure 2H and 2I**). In addition, the percentages of T-bet^+^CD4^+^, ROR-γt^+^CD4^+^ and Foxp3^+^CD4^+^ T cells in spleen, MLN and colonic LPL (**Figure 2J and 2K**), and IFN-γ^+^CD4^+^ and IL-17A^+^CD4^+^ T cells in spleen and MLN (**Figure S4B**) showed no significant difference between ABX+DSS+ group and ABX+DSS+CSR+ group. These results demonstrated that the protective effect of CSR on colitis was gut microbiota-dependent.

**Figure 2.**
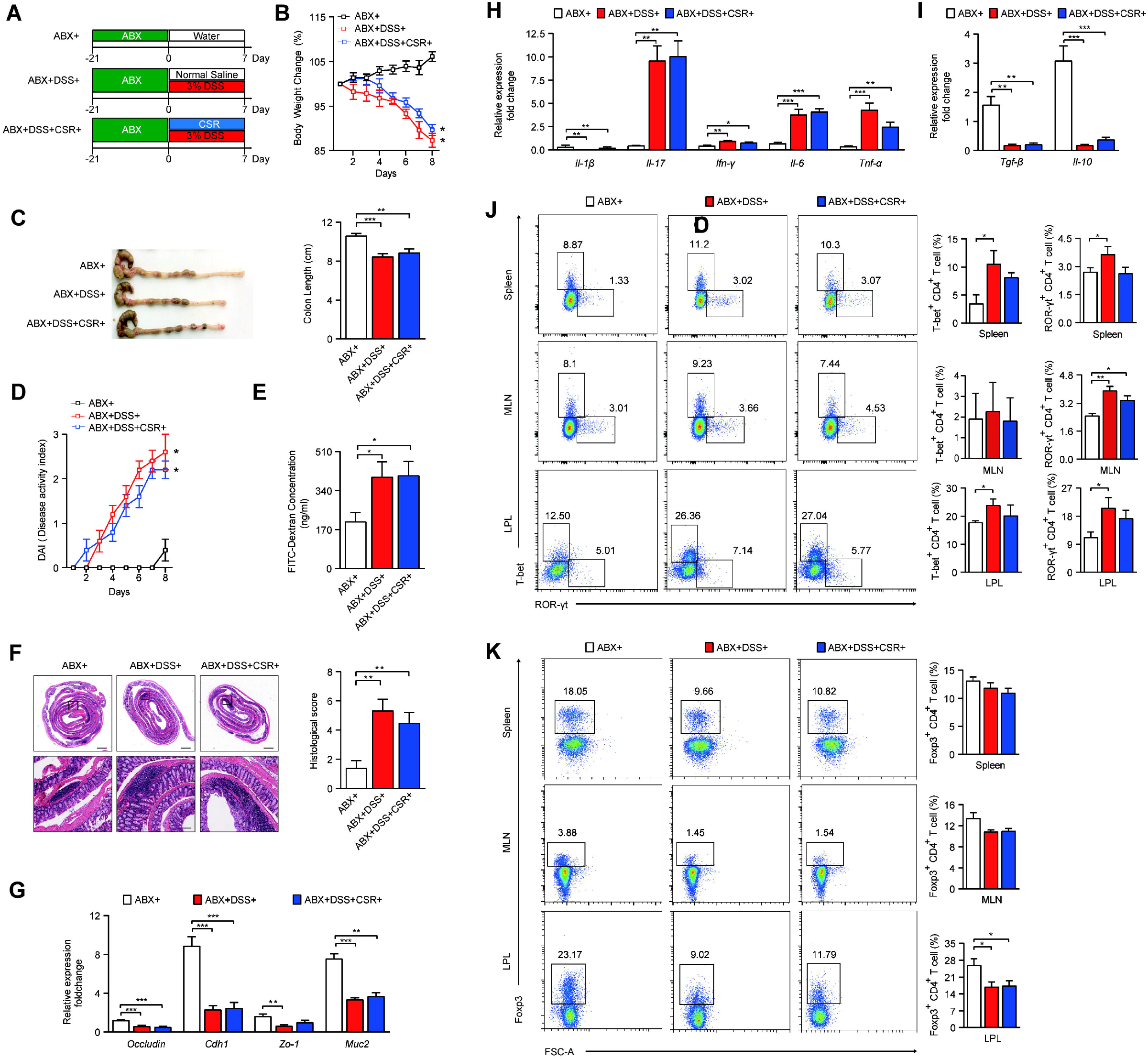
Effects of CSR against DSS-induced colitis after pretreatment with ABX. (A) Schematic diagram illustrates the experimental design. (B) Body weight percentage changes of each group. (C) Measurement of the length of colons harvested from mice in each group (n=10). (D) The effect of CSR on DAI in mice (n=10). (E) Mice in each group received an oral gavage of FITC-dextran (0.5mg/g) and serum FITC-dextran concentrations were determined 4□h later (n=5). (F) H&E staining (Bar=1mm above, Bar=100um below) sections and histological scores of colon tissue from mice in each group (n=5). (G) The expression levels of Occludin, Cdh1, ZO-1 and MUC2 of colon tissue in each group (n=5). (H) The expression levels of pro-inflammatory cytokines of colon tissue in each group (n=5). (I) The expression levels of anti-inflammatory cytokines of colon tissue in each group (n=5). (J) T-bet^+^CD4^+^ (Th1) cells and ROR-γt^+^CD4^+^ (Th17) cells in spleen, MLN and LPL from the ABX+ group, ABX+DSS+ group and ABX+DSS+CSR+ group were analyzed by flow cytometry and bar charts of the percentage of Th1 and Th17 cells (n=5). (K) Foxp3^+^CD4^+^ (Treg) cells in spleen, MLN and LPL from ABX+ group, ABX+DSS+ group and ABX+DSS+CSR+ group were analyzed by flow cytometry and bar charts of the percentage of Treg cells (n=5). Data are presented as mean ± *SEM*. \**P* < 0.05, significantly different as indicated.

### Fecal microbial transplantation mitigated colitis

To gain deeper insight into the protective effect of CSR about how it regulated the gut microbiota, we conducted FMT experiment. Fecal microbiota from mice of the DSS-CSR-, DSS+CSR- and DSS+CSR+ group were transferred into DSS-induced colitis mice (**Figure 3A**). Compared to mice with FMT from DSS-CSR-treated mice, mice received FMT from DSS+CSR-treated mice showed more serious weight loss (**Figure 3B**) and colon shortening (**Figure 3C**), higher DAI score (**Figure 3D**), poorer intestinal permeability (**Figure 3E**), higher histology score (**Figure 3F**), lower levels of goblet cells (**Figure S5A**) and tight junction proteins (**Figure 3G**), which were completely reversed in mice with FMT from DSS+CSR+ treated mice. Representative microscopic H&E staining and Alcian blue staining pictures are shown in **Figure 3F** and **Figure S5A**. Furthermore, compared to mice with FMT from DSS-CSR-treated mice, the expression of related proinflammatory cytokines, Th1, Th17 cells and their transcription factors were increased, and the anti-inflammatory counterparts were decreased in mice received fecal microbiota from DSS+CSR-mice. Again, all aforementioned disease-modifying tendencies were abrogated in mice received fecal microbiota from DSS+CSR+ mice (**Figure 3H and 3K, Figure S5B**). These FMT results indicated that gut microbiota in CSR treated mice was responsible for alleviated colitis.

**Figure 3.**
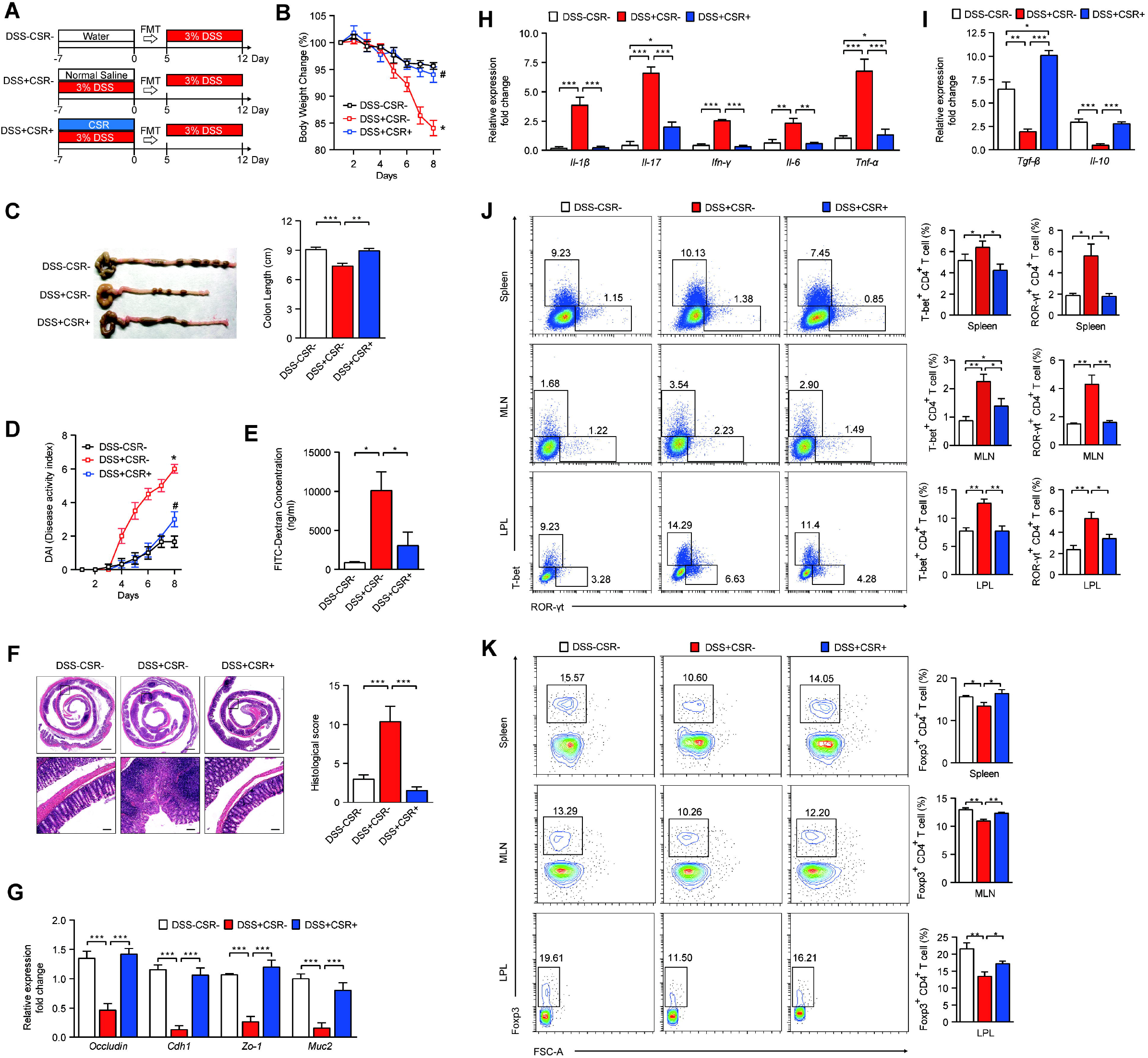
Fecal transplants from CSR-treated mice confer the protection for colitic mice. (A) Schematic diagram illustrates the experimental design. (B) Body weight percentage changes of each group. (C) Measurement of the length of colons harvested from mice in each group (n=10). (D) The effect of CSR on DAI in mice (n=10). (E) Mice in each group received an oral gavage of FITC-dextran (0.5mg/g) and serum FITC-dextran concentrations were determined 4□h later (n=5). (F) H&E staining (Bar=1mm above, Bar=100um below) sections and histological scores of colon tissue from mice in each group (n=5). (G) The expression levels of Occludin, Cdh1, ZO-1 and MUC2 of colon tissue in each group (n=5). (H) The expression levels of pro-inflammatory cytokines of colon tissue in each group (n=5). (I) The expression levels of anti-inflammatory cytokines of colon tissue in each group (n=5). (J) T-bet^+^CD4^+^ (Th1) cells and ROR-γt^+^CD4^+^ (Th17) cells in spleen, MLN and LPL from DSS-CSR-group, DSS+CSR-group and DSS+CSR+ group were analyzed by flow cytometry and bar charts of the percentage of Th1 and Th17 cells (n=5). (K) Foxp3^+^CD4^+^ (Treg) cells in spleen, MLN and LPL from the DSS-CSR-group, DSS+CSR-group and DSS+CSR+ group were analyzed by flow cytometry and bar charts of the percentage of Treg cells (n=5). Data are presented as mean ± *SEM*. \**P* < 0.05, significantly different as indicated.

### Celastrol significantly influenced the gut microbiota

The gut microbiota has been shown to play important roles in maintaining intestinal homeostasis and T cell functions. High-throughput gene-sequencing analysis of 16S rRNA in fecal bacterial DNA isolated from control group, DSS+ group and DSS+CSR+ group mice were performed to investigate whether CSR altered microbiome. We initially measured gut microbial alpha diversity through different indices, including observed species, Chao, ace and PD_whole tree, and found that CSR-treated mice harbored a microbiota with higher alpha diversity relative to that of the DSS+ group (**Figure 4A**). To further explore the diversity of the microbiome, we performed beta-diversity to generate principal coordinate analysis (PCoA) using binary jaccard distance and unweighted-unifrac distance algorithms. The obvious clustering separation between OTUs reveals the different community structures of the three groups, indicating that these communities are different in composition and structure (**Figure 4B**).

**Figure 4.**
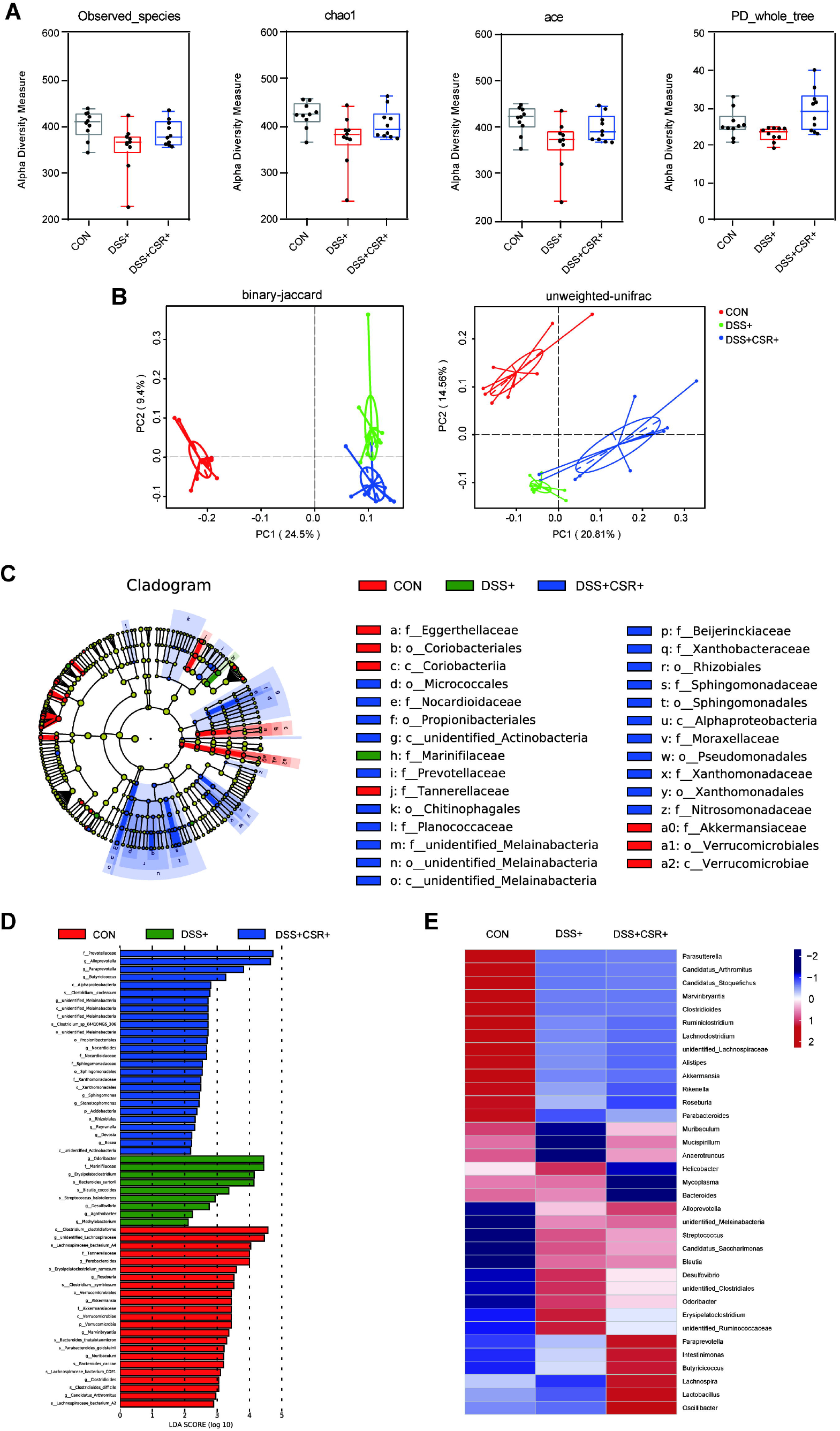
CSR treatment significantly altered the gut microbiota diversity and composition. (A) Alpha diversity boxplot (observed species, Chao1, ace and PD-whole tree). (B) Principal coordinate analysis (PCoA) using Binary-Jaccard and Unweighted-UniFrad of beta diversity. (C) Taxonomic cladogram from LEfSe, depicting taxonomic association from between microbiome communities from the control, DSS+ and DSS+CSR+ groups. (D) LDA score computed from features differentially abundant between the control, DSS+ and DSS+CSR+ groups. (E) Heatmap of selected most differentially abundant features at the genus level. The blue color represents less abundant, white color represents intermediate abundance and red represents the most abundant. Each symbol represents an individual mouse.

Subsequently, we evaluated the gut microbiota in all samples to find possible compositional differences among the three groups. At the phylum level, in all samples the most abundant phyla were *Bacteroidetes* and *Firmicutes*. Additionally, the ratio of *Firmicutes* to *Bacteroidetes* (F/B) showed no significant difference among the three groups **(Figure S6A)**. Taxonomic compositions of the three groups were also compared at the class/order/family level **(Figure S6B and S6D)**. At the genus level, the three groups displayed obviously differential biological compositions **(Figure S6E)**. The different bacterial genera with higher relative abundance in three groups were analyzed. *Alloprevotella* and *Odoribacter* displayed relatively higher abundance in DSS+ and DSS+CSR+ groups, whereas the *unidentified_Lachnospiraceae* were more abundant in control group **(Figure S6F)**.

To confirm which bacteria was changed by CSR treatment, we conducted high-dimensional class comparisons using the linear discriminant analysis (LDA) of effect size (LEfSe), and looked for differences in the predominance of bacterial communities among the three groups (**Figure 4C and 4D**). According to the analysis results, *Odoribacter* and *Marinifilaceae* were the key types of bacteria resulting in the gut microbiota dysbiosis in DSS-treated group. However, *Prevotellaceae, Alloprevotella, Paraprevotella* and *Butyricicoccus* displayed a relative enrichment in the DSS+CSR+ group, which might be associated with the CSR-mediated alleviation of colitis. Based on the OUT abundance at genus levels, the inter-group difference in the comparison heatmap was shown for analyzing gut microbiota among the three groups (**Figure 4E**). Similarly, the genus *Alloprevotella* and *Butyricicoccus* displayed a relatively higher abundance in the DSS+CSR+ group, while the genus *Odoribacter* was significantly enriched in the DSS+ group, which was consistent with the LEfSe analysis results. In conclusion, CSR treatment significantly altered the gut microbiota diversity and composition.

### Celastrol significantly alters gut metabolomics

Metabolites were the major executors of gut microbiota in the pathogenesis of inflammatory diseases. Hence, to address the role of CSR in affecting gut metabolites, we performed non-target metabolomics analysis on the isolated feces of mice from DSS+ group and DSS+CSR+ group. Partial least-squares discrimination analysis (PLS-DA) and Hierarchical clustering revealed that the metabolomic profile of DSS+CSR+ mice was significantly different from that of DSS+ mice (**Figure 5A and 5B**). Under negative and positive ion mode, 24 metabolites were more abundant in DSS+CSR+ mice than in DSS+ mice (**Figure 5C and 5D**). Then the influences of these metabolites on the differentiation of naïve CD4+ T cells were explored in vitro, and pyruvate and adenosine showed their potential functions. In details, pyruvate down-regulated the differentiation of IFN-γ^+^CD4^+^ T cells and IL-17A^+^CD4^+^ T cells (**Figure 6A and 6B**), and adenosine up-regulated the differentiation of Foxp3^+^CD4^+^ T cells in vitro (**Figure 6C**). In summary, CSR administration obviously altered the metabolomics in colon, which in turn affected the differentiation of naïve T cells.

**Figure 5.**
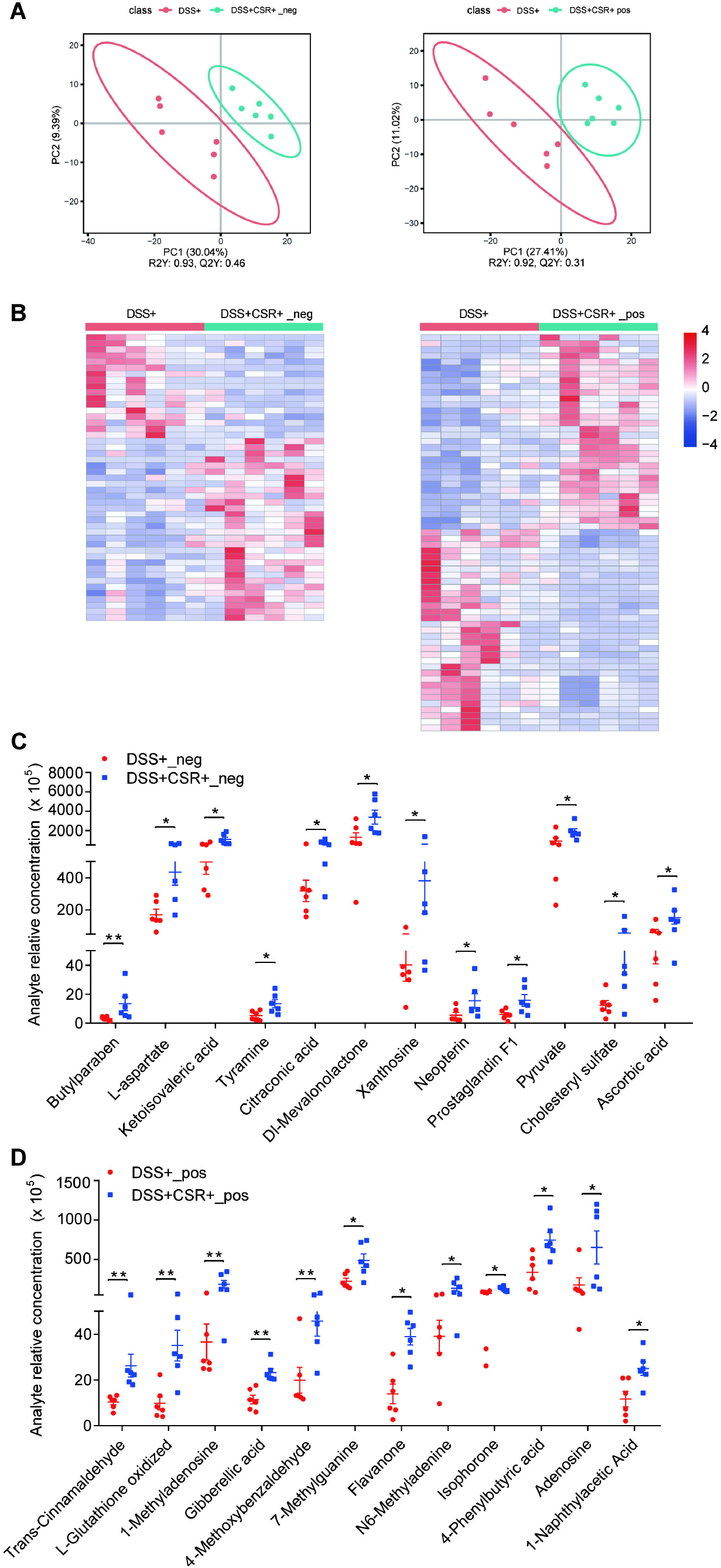
CSR treatment significantly alter metabolomics. (A) Partial least-squares discrimination analysis (PLS-DA) of metabolomic profile. (B) Hierarchical clustering of metabolites. (C) Relative concentration of metabolites under negative ion mode in DSS+ group and DSS+CSR+ group. (D) Relative concentration of metabolites under positive ion in DSS+ group and DSS+CSR+ group. Data are pooled in one independent experiment and presented as mean ± *SEM*. \**P* < 0.05, significantly different as indicated.

**Figure 6.**
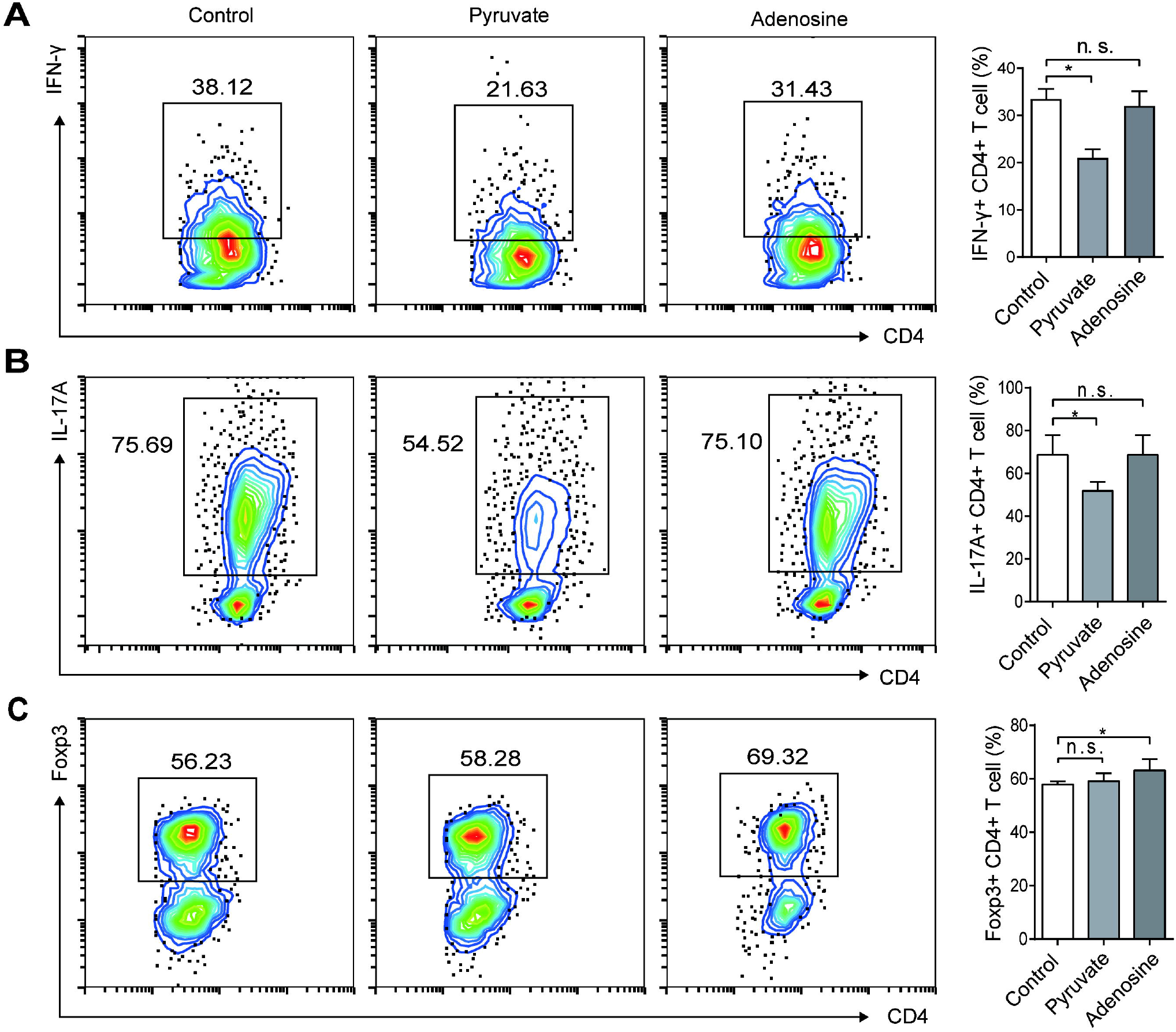
The influence of pyruate and adenosine treatment on T cell differentiation in vitro. Spleen naive CD4^+^ T cells from C57BL/6 were cultured under Th1, Th17 and Treg skewing condition in the presence or absence of pyruate or adenosine respectively. (A) A plot from one representative experiment displays the proportions of IFN-γ^+^CD4^+^ (Th1) cells, and the mean proportions of Th1 cells were analyzed by bar chart. (B) A plot from one representative experiment displays the proportions of IL-17A^+^CD4^+^ (Th17) cells, and the mean proportions of Th17 cells were analyzed by bar chart. (C) A plot from one representative experiment displays the proportions of Foxp3^+^CD4^+^ (Treg) cells, and the mean proportions of Treg cells were analyzed by bar chart. Data are presented as mean ± *SEM*. \**P* < 0.05, significantly different as indicated.

## Discussion

UC is a subtype of inflammatory bowel disease (IBD) characterized by chronic recurrent inflammation of the colonic mucosa. The clinical manifestations are recurrent abdominal pain, loose and bloody stools. Currently, drugs applied in the treatment are challenged by insufficient effect, drug dependence, adverse reactions and high costs. Thus, it is worthy of in-depth investigation to find new UC therapies with better performance. CSR, with prominent anti-inflammatory and antioxidant effects, is anticipated to be an effective element in TW, which has been “clinically” tested in the Traditional Chinese Medicine for thousand years. In this study, we systematically studied the therapeutic effects of CSR on DSS-induced colitis in mice and its potential mechanism (**Figure 7**).

**Figure 7.**
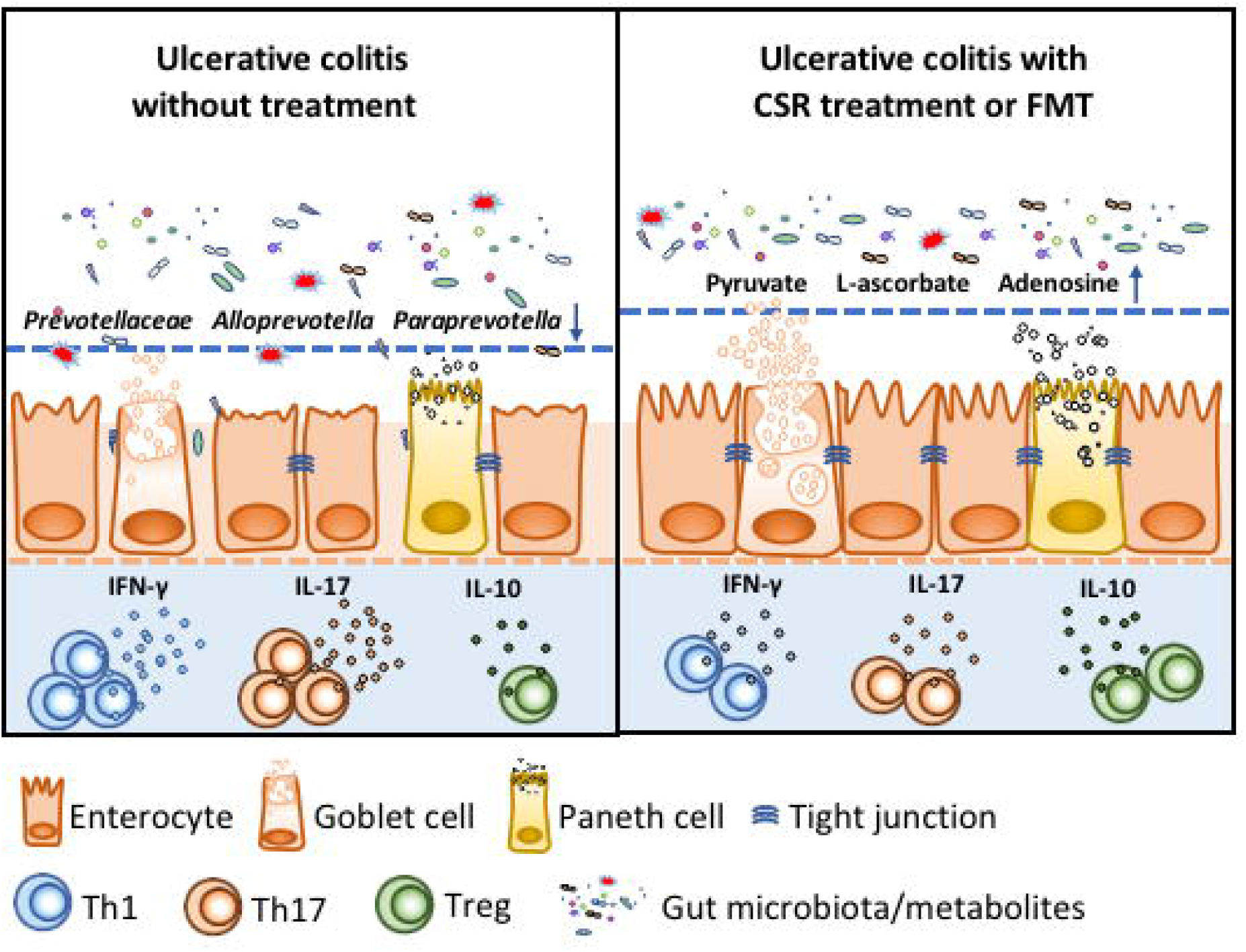
Schematic depiction about the protective effects of celastrol during ulcerative colitis.

The symptom of DSS-induced colitis in mice is similar to that of human UC, and the DSS-induced colitis model is well-acknowledged and wide-used animal model for studying UC currently[24]. DSS-induced colitis in mice reveals the typical UC features, including inflammation that starts from the distal colon and then involves the proximal colon, body weight loss, shortening of colon length, mucosal ulcers, and infiltration of inflammatory granulocytes[25]. According to the results, CSR significantly affected the intensity of intestinal inflammation, reversing the imbalance of Treg/Th17 and Treg/Th1 in the intestinal mucosa. These results are similar to those reported in previous studies, where the beneficial effects of CSR on the Th17/Treg cell-mediated responses were revealed in animal models of autoimmune arthritis and autoimmune encephalomyelitis[26, 27].

Gut microbiota dysbiosis plays a vital role in the pathogenesis of UC[28, 29], and various therapeutic microbial manipulations (such as antibiotics, probiotics, prebiotics, and microbiota transplantation) have been proved to be a promising treatment strategy[30–32]. To reveal the underlying therapeutic mechanism of CSR action, we investigated whether gut microbiota contributes to its protective effect. It is worth noting that there was no significant difference in the severity of colon inflammation between the ABX+DSS+ group and the ABX+DSS+CSR+ group, which means that the protective role of CSR disappeared after depleting gut microbiota. Subsequently, we conducted FMT to confirm that the effect was afforded by microbiota and was transferable. In contrast to the feces from DSS+CSR-mice, feces from DSS+CSR+ mice alleviated the inflammatory response and rectified the imbalance of Treg/Th1 and Treg/Th17 in DSS-induced mice. The experiments in vitro have confirmed that, without the participation of microbiota, CSR indeed lost its regulatory role in the differentiation of Th1, Th17 cells, and Treg cells. Therefore, the immune-mediating action of CSR was due to its influence on the microbiota.

According to the literature, UC patients show a decrease in the biological diversity of gut microbiota composition, which is called dysbiosis and characterized by the loss of beneficial bacteria and the expansion of pathogenic bacteria[14]. For instance, in UC patients, the relative abundance of beneficial bacterial species, such as *Prevotella copri* and the butyrate-producing bacterium *Faecalibacterium prauznitzii*, have been shown to decrease remarkably. In order to further clarify the influence of CSR on the structure and composition profiles of gut microbiota, the 16S rRNA sequencing was conducted. In this study, the alpha diversity indices, including observed species, Chao, ace and PD_ whole tree indexes, revealed that CSR-treated mice were characterized by a higher diversity relative to that of the DSS-treated group. The beta diversity analysis has revealed the DSS+CSR+ group mice harbored an apparent clustering separation from DSS+ group mice through PCoA, indicating that CSR treatment markedly transformed the biological community structures. Consequently, the conducted LEfSe analysis between the control, DSS+, and DSS+CSR+ groups revealed that *Odoribacter* and *Marinifilaceae* were the key types of bacteria in DSS-treated group. Meanwhile, four signature bacterial taxa, including *Prevotellaceae*, *Alloprevotella*, *Paraprevotella* and *Butyricicoccus* displayed a relative enrichment in the DSS+CSR+ group. Given the dysbiosis in IBD patients is associated with a decrease in the number of SCFAs/butyrate-producing bacteria[33], the obtained results may indicate a favorable action of CSR on the course of UC. Moreover, the genus *Prevotellaceae*, with the most predominance and the highest LDA score in the DSS+CSR+ group, was reported to be associated with the remission of IBD[34]. Additionally, *Alloprevotella* was reported to be related to the decreased lifetime of cardiovascular disease[35], which strengthen the importance of the present findings.

Previous study illuminated that the metabolites of gut microbiota, such as SCFAs, impose profound impact on inflammation of IBD[36]. We used untargeted metabolomics analysis to investigate the potential alteration in metabolome after CSR treatment. The results showed that the concentration of pyruvate, L-ascorbate and adenosine were higher in DSS+CSR+ group mice. Pyruvate is a key metabolite of microbial cells, and the end product of glycolysis as well as the major metabolite of amino acid and protein metabolism. The acid could prevent hydrogen peroxide-induced apoptosis and enhance the metabolism of fatty acid[37, 38]. Fatty acid, with the effects of anti-inflammation and promoting autophagy, is obviously decreased in patients with UC. Moreover, indole-3-pyruvate was also proved to alleviate the inflammation of colon in experimental colitis mice[39]. L-ascorbate could attenuate endotoxin-induced production of inflammatory mediators by inhibiting MAPK activation and NF-κB translocation, the two signal pathways are activated in IBD[40]. It has also been reported that coordination of ENT2-dependent adenosine transport and signaling could dampen mucosal inflammation[41]. All in all, the upregulated pyruvate, L-ascorbate and adenosine might perform their anti-inflammatory effect in certain way in colitis.

In conclusion, our data demonstrated that CSR ameliorated colon inflammation in a gut microbiota-dependent manner. While the anti-inflammatory and immunosuppressive properties of CSR have been well described and discussed[21], its action on the gut microbiota was not studied. Thus, we can conclude that the underlying protective mechanism of therapeutic action of CSR is associated not only with rectifying the Treg/Th1 and Treg/Th17 balances and downregulation of inflammatory cytokine but with the modulation of microbiota-community structure and metabolisms as well. While the exact role of microbiome needs to be further investigated, this study still opens up a new direction in the study of traditional medicines.

## Methods

### Animals

All animal care and experimental procedures were approved by the Committee for Animal Research of Huazhong University of Science and Technology (Wuhan, China). Male BALB/c mice (8 weeks old, 18-22 g) were purchased from the experimental animal center of Huazhong University of Science and Technology (Wuhan, China). These animals were socially housed at relatively constant humidity (40-60%), temperature (22-24°C) and a 12-hours light/dark cycle and maintained on a normal chow diet with free access to water. All mice were allowed for 1 week housing before the experiment, and then randomly separated into three groups: normal control, DSS+, and DSS+CSR+. The acute experimental colitis model was induced according to previous publication[42]. Briefly, mice were administrated with 3.0% (wt/vol) dextran sodium sulfate (DSS, MP Biomedicals, UK) supplemented in filter-purified drinking water for consecutive 7 days. After adaptive feeding, the DSS+CSR+ and DSS+ groups were performed for colitis induction, and administrated simultaneously with CSR (Chem Faces, Cat# CFN99198) at a dose of 1 mg·kg^-1^ and equal amount of saline by gavage daily for 7 days, respectively. Similarly, the normal control group were given distilled water with equal amounts of saline by gavage. During DSS treatment, the morbidity, body weights, stool consistency and stool occult blood of mice were daily monitored. The severity of colitis was measured by disease activity index (DAI) as described before[43]. The colon length was measured when the mice were were euthanized by excessive pentobarbital sodium.

### Depletion of the gut microbiota

For the gut microbiota depletion experiment, mice were randomly divided into three groups: ABX+, ABX+DSS+, and ABX+DSS+CSR+. Mice in these three groups were treated with an antibiotic cocktail, including 1 g·L^-1^ ampicillin (Macklin), 1 g·L^-1^ neomycin (Sigma), 1 g·L^-1^ metronidazole (Sigma) and 0.5 g·L^-1^ vancomycin (Macklin), in drinking water for 3 weeks. Subsequently, the mice in ABX+DSS+ and ABX+DSS+CSR+ groups were provided with DSS in drinking water, and mice in the ABX+DSS+CSR+ group were treated with CSR (1 mg·kg^-1^) orally once daily during DSS treatment whereas mice in other groups were treated with equal amount of saline.

### Fecal microbiota transplantation

The fecal microbiota transplantation was performed based on the protocol as described before[44]. Briefly, donor mice were randomly divided into three groups including the DSS-CSR-, DSS+CSR-, and DSS+CSR+ groups. The DSS+CSR- and DSS+CSR+ groups received DSS diluted in drinking water to induce colitis. The DSS+CSR+ group received CSR treatment (1 mg·kg^-1^) orally once per day whereas the DSS-CSR- and DSS+CSR-groups received saline. After 7 days, the stools from each donor group were collected daily under a laminar flow hood in sterile conditions. Then the samples were pooled, and 100 mg was resuspended in 1 ml of sterile saline.

The solution was vigorously mixed for 10 s followed by centrifugation at 800 g for 3 min. Then the supernatant was collected and used as transplant material within 10 min by oral gavage to prevent changes in bacterial composition. Recipient mice were randomly divided into three groups. Each group of recipient mice received DSS treatment to induce colitis and simultaneously 200 ul freshly prepared supernatant above per day for 7 days.

### Intestinal permeability assays

Intestinal permeability was assessed by the fluorescein isothiocyanate conjugated dextran (FITC-dextran) tracer (4 kDa, Sigma-Aldrich, Cat# 68059). Mice were fasted overnight and then administered with 0.5 ml of FITC-dextran (0.5 mg·g^-1^ body weight) by oral gavage 4 h before being euthanized. Blood samples were collected at the time of euthanization, and the samples were centrifuged for 90 s at 6000 g. FITC-dextran concentration was determined using fluorescence spectrometry at an excitation wavelength of 488 nm and emission wavelength of 520 nm within 5 min[45].

### Histopathology and Alcian blue staining

The colons were emptied of fecal contents and opened longitudinally along the mesenteric border and formed a Swiss-Roll from the proximal to the distal end, then placed in 4% paraformaldehyde for 24 hours. The Swiss-Rolls were transferred to paraffin-embedded blocks to generate 5-μm-thick sections for hematoxylin and eosin (H&E) staining and assessed blindly by a pathologist. The 5-μm-thick sections were also stained with Alcian blue periodic acid by standard techniques. Slices were visualized under a light microscope (Leica, Germany).

### Mouse colonic lamina propria lymphocytes (LPL) isolation

Mouse colons were opened longitudinally and washed with cold PBS with 1M Hepes to remove the fecal contents. Pooled colons were cut into 0.5 cm pieces and washed with 25 ml of HBSS (Gibco, Cat# C14175500BT) containing 1M Hepes and 500 mM EDTA on an orbital shaker at 100 rpm for 25 min at 37 °C. Then the colon pieces were washed twice using cold PBS with 1 M Hepes. After washing, the colons were finely cut and digested with 10 ml RPMI 1640 (Gibco, Cat# 11875-119) containing 1mg·ml^-1^ DNase I (Roche, Cat# 10104159001), 0.5 mg·ml^-1^ Type-D Collagenase (Roche, Cat# 11088858001) at 100 rpm for 15 min at 37 °C. After digestion, the colonic LP cells were filtered through 100 μm strainer, followed by centrifugation at 1650 rpm for 5 min at 4 °C, and resuspended on 500 μl PBS for flow cytometric analysis[46].

### Flow cytometry

Flow cytometry was performed as described previously[47]. Briefly, Single cell suspensions were stained with indicated antibodies diluted by PBS supplemented with 2% FBS and 0.5% BSA for surfaces markers. For the staining of intracellular cytokines IFN-γ and IL-17A, cells were incubated and stimulated with 200 ng·ml^-1^ phorbol myristate acetate (PMA) (Enzo, Cat# BML-PE160-0005), 1 μg·ml^-1^ ionomycin (Enzo, Cat# ALX-450-007-M001), 1 μg·ml^-1^ brefeldin A (eBioscience, Cat# 00-4506-51) at 37 °C for 6 h. Then, cultured cells were collected, washed and stained surface markers with CD45 and CD4 for 20 min, followed by fixation and permeabilization, and then stained intracellularly with anti-IFN-γ and anti-IL-17A antibody for 30 min. For Foxp3, T-bet and ROR-γt staining, the cells were stained for surface marker such as CD45 and CD4, followed by fixation and permeabilization with fixation and permeabilization buffer (Thermo Fisher Scientific, Cat# 88-8824-00) at room temperature for 30 min. After washes, the cells were then stained with anti-Foxp3, anti-T-bet or anti-ROR-γt antibody as instructed. All samples were detected by CytoFLEX LX Flow Cytometry System and analyzed with the CytExpert 2.0 Software. Antibodies used for flow cytometry included anti-mouse CD45-FITC (BD Biosciences, Cat# 553080), CD4-PE/Cy7 (BD Biosciences, Cat# 552775), IL-17A-PE (BD Biosciences, Cat# 561020), IFN-γ-APC (BD Biosciences, Cat# 562018), ROR-γt-BV421 (BD Biosciences, Cat# 562894), Foxp3-PE (Thermo Fisher Scientific, Cat# 12-5773-82), T-bet-APC (BD Biosciences, Cat# 561264), and Fixable Viability Stain 510 (BD Biosciences, Cat# 564406). Flow gating strategies for each cell population were showed in **Figure S7**.

### Quantitative real-time polymerase chain reaction (qRT-PCR) for mRNA

Total RNA was extracted from colonic tissues using RNAiso Plus (TaKaRa, Cat# 9109) according to the manufacturer’s protocols. After isolation of RNA, PrimeScript RT Master Mix (TaKaRa, Cat# RR036A) were used to generate complementary DNAs (cDNAs) of mRNAs. Then, these cDNAs were analyzed to explore genes expression changes by SYBR Premix Ex Taq (TaKaRa, Cat# RR420A). The relative expression levels of genes in tissues were normalized to β-actin. The sequences of all primers are listed in **Table 1**.

**Table 1.**
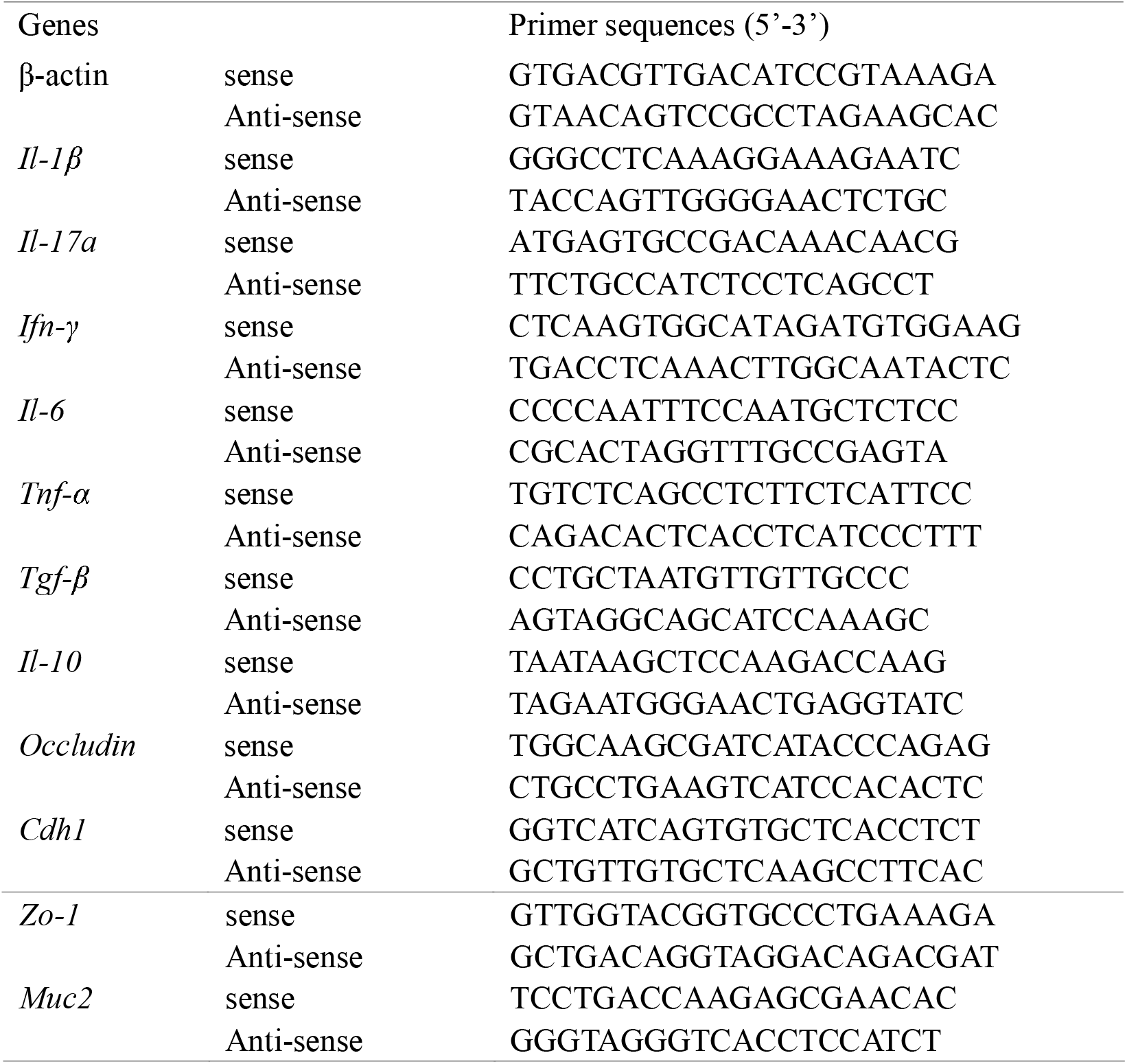
Primer sequences of target genes for mice.

### Fecal genomic DNA extraction and 16S-rRNA sequencing

Fecal genomic DNA extraction and 16S-rRNA sequencing were performed at Novogene (Beijing, China). Total genome DNA from fecal samples was extracted using CTAB/SDS method. DNA concentration and purity were monitored on 1% agarose gels. According to the concentration, DNA was diluted into 1 ng·μl^-1^ using sterile water. 16S rRNA genes of distinct regions (16S V3-V4) were amplified used specific primer with the barcode. All PCR reactions were carried out with Phusion^®^ High-Fidelity PCR Master Mix (New England Biolabs). Mix same volume of 1×loading buffer (contained SYBR green) with PCR products and operate electrophoresis on 2% agarose gel for detection. PCR products was mixed in equidensity ratios. Then, mixture PCR products was purified with GeneJET^™^ Gel Extraction Kit (Thermo Fisher Scientific). Sequencing libraries were generated using Ion Plus Fragment Library Kit 48 rxns (Thermo Fisher Scientific) following manufacturer’s recommendations. The library quality was assessed on the Qubit@ 2.0 Fluorometer (Thermo Fisher Scientific). At last, the library was sequenced on an Ion S5^™^ XL platform and 600 bp single-end reads were generated.

### Untargeted metabolomics analysis

Untargeted metabolomics analysis were performed at Novogene (Beijing, China). Fecal samples (100 mg) were individually grounded with liquid nitrogen and the homogenate was resuspended with prechilled 80% methanol and 0.1% formic acid by fully vortexing. The samples were incubated on ice for 5 min and then were centrifuged at 15000 rpm, 4°C for 5 min. And then, the supernatant was diluted to final concentration containing 53% methanol by LC-MS grade water. The samples were subsequently transferred to a fresh Eppendorf tube and then were centrifuged at 15000 g, 4°C for 10 min. Finally, the supernatant was injected into the LC-MS system analysis.

### Isolation of naïve CD4^+^ T cells and in vitro induction of differentiation

Total spleen T cells were purified by negative selection with the mouse naïve CD4^+^ T Cells Isolation Kit (Miltenyi Biotec, Cat# 130-104-453) and LS separation columns (Miltenyi Biotec, Cat# 130-042-401) following the manufacturer’s instructions. Naive CD4^+^ T cells were seeded in 96-well plates coated with anti-CD3 (Thermo Fisher Scientific, Cat# 16-0032-85) and anti-CD28 (Thermo Fisher Scientific, Cat# 16-0281-85) antibodies at 2 x 10^5^ cells/well in RPMI 1640 medium (Gibco, Cat# 11875-119) containing 10% inactivated fetal bovine serum (Gibco, Cat# 10099141). For Th1 cell differentiation, naïve CD4^+^ T cells were cultured with 10 ng·ml^-1^ IL-12 (R&D, Cat# 402-ML-020/CF) and 10 μg·ml^-1^ anti-IL-4 (Bio X Cell, Cat# BE0045, RRID:AB_1107707). For Th17 cell differentiation, naive CD4^+^ T cells were cultured with 2 ng·ml^-1^ TGF-β (R&D, Cat# 240-B-002/CF), 10 ng·ml^-1^ IL-6 (R&D, Cat# 406-ML-005/CF), 10 μg·ml^-1^ anti-IFN-γ (Bio X Cell, Cat# BE0055, RRID:AB_1107694), 10 μg·ml^-1^ anti-IL-4 and 10 μg·ml^-1^ anti-IL-2 (Bio X Cell, Cat# BE0043-1, RRID:AB_1107705). For Treg cell differentiation, naive CD4^+^ T cells were cultured with 2 ng·ml^-1^ TGF-β.

The effects of different concentrations of CSR, pyruate (Macklin, Cat# P6033) and adenosine (Macklin, Cat# A6218) on the differentiation of Th1, Th17 and Treg were assessed by flow cytometry after 72 h culture.

### Data and analysis

All experiments were designed to generate groups of equal size, using randomization and blind data analysis, and no data points were excluded from the analysis in any experiment. The data of qRT-PCR were normalized to control to avoid unwanted sources of variation. The statistical analysis was undertaken only for studies where each group size was at least n=5. Data are presented as mean ± *SEM*, and statistic were analysed with GraphPad Prism (GraphPad Prism version 6.0). Student’s *t*-tests were used to compare the means of two groups. And comparisons among multiple groups were performed with one-way ANOVA with Tukey’s test. Post hoc tests were run only if F achieved *P* < 0.05, and there was no significant variance inhomogeneity. *P*-values < 0.05 were considered statistically significant. The declare group size is the number of independent values, and statistical analysis was performed using these independent values (technical replicates were not treated as independent values).

## Abberviations

UC: ulcerative colitis
CSR: celastrol
FMT: fecal microbiota transplantation
Th cells: T helper cells
Treg: regulatory T
DSS: dextran sodium sulfate
LPL: lamina propria lymphocytes.

## Author contribution

Fan Heng: Project administration. Li Mingyue, Tian Chunxia, Gui Yang and Dong Yalan: Conceptualization, Formal analysis, Methodology, Investigation, Software, Writing-original draft. Yu Ting, Guo Weina, Wang Wenzhu and Zhang Zili: Validation. Xue Kaming and Jiang Feng: Date curation. Li Junyi, Zhou Haifeng and Alexey Sarapultsev: Methodology. Hu Desheng and Luo Shanshan: Supervision, Funding acquisition.

## Acknowledgments

This study was funded by the grants from the National Key R&D Program of China (2019YFC1316204), and the National Natural Science Foundation of China (Nos. 81974249, 31770983, 82070136), and the Hubei Provincial Natural Science Foundation of China (No.2020BHB016).

## Supplementary material

**Figure S1. CSR attenuated DSS-induced experimental colitis in mice.**

(A) The molecular structure of CSR. (B) Alcian blue staining (Bar=100um) sections of colon tissue from mice in each group (n=5). (C) IFN-γ^+^CD4^+^ (Th1) cells and IL-17A^+^CD4^+^ (Th17) cells in spleen and MLN from the control, DSS+ group and DSS+CSR+ group were analyzed by flow cytometry and bar charts of the percentage of Th1 and Th17 cells (n=5). Data are presented as mean ± *SEM*. \**P* < 0.05, significantly different as indicated.

**Figure S2. The influence of celastrol treatment on T cell differentiation in vitro.**

Spleen naive CD4^+^ T cells from C57BL/6 were cultured under Th1, Th17 and Treg skewing condition in the presence or absence of CSR respectively. (A) A plot from one representative experiment displays the proportions of IFN-γ^+^CD4^+^ (Th1) cells, and the mean proportions of Th1 cells were analyzed by bar chart. (B) A plot from one representative experiment displays the proportions of IL-17A^+^CD4^+^ (Th17) cells, and the mean proportions of Th17 cells were analyzed by bar chart. (C) A plot from one representative experiment displays the proportions of Foxp3^+^CD4^+^ (Treg) cells, and the mean proportions of Treg cells were analyzed by bar chart. Data are presented as mean ± *SEM*. \**P* < 0.05, significantly different as indicated.

**Figure S3. General changes of mice after antibiotics treatment.**

(A) ABX treatment altered the total DNA and the structure of gut microbiota in fecal samples of mice (n=5). (B) H&E staining indicated the antibiotics treatment could not affecte the morphology of liver (Bar=50um), kidney (Bar=50um), and colon (Bar=100um) (n=5); (C) Serum ALT, AST, BUN and CRE levels were not changed after antibiotics treatment (n=5). Data are presented as mean ± *SEM*. \**P* < 0.05, significantly different as indicated.

**Figure S4. Effects of CSR against DSS-induced colitis after pretreatment with ABX.**

(A) Alcian blue staining (Bar=100um) sections of colon tissue from mice in each group (n=5). (B) IFN-γ^+^CD4^+^ (Th1) cells and IL-17A^+^CD4^+^ (Th17) cells in spleen and MLN from the ABX+ group, ABX+DSS+ group and ABX+DSS+CSR+ group were analyzed by flow cytometry and bar charts of the percentage of Th1 and Th17 cells (n=5). Data are presented as mean ± *SEM*. \**P* < 0.05, significantly different as indicated.

**Figure S5. Fecal transplants from CSR-treated mice confer the protection for colitic mice.**

(A) Alcian blue staining (Bar=100um) sections of colon tissue from mice in each group (n=5). (B) IFN-γ^+^CD4^+^ (Th1) cells and IL-17A^+^CD4^+^ (Th17) cells in spleen and MLN from DSS-CSR-group, DSS+CSR-group and DSS+CSR+ group were analyzed by flow cytometry and bar charts of the percentage of Th1 and Th17 cells (n=5). Data are presented as mean ± *SEM*. \**P* < 0.05, significantly different as indicated.

**Figure S6. CSR treatment significantly altered the gut microbiota diversity and composition**.

(A) Bar plots of the phylum taxonomic levels in control, DSS+ group and DSS+CSR+ group. Relative abundance is plotted for each sample. The relative abundances of Bacteroidetes and Firmicutes, and the ratio of Firmicutes to Bacteroidetes (F/B). (B) Bar plots of the class taxonomic levels in control, DSS+ group and DSS+CSR+ group. Relative abundance is plotted for each sample. (C) Bar plots of the order taxonomic levels in control, DSS+ group and DSS+CSR+ group. Relative abundance is plottec for each sample. (D) Bar plots of the family taxonomic levels in control, DSS+ group and DSS+CSR+ group. Relative abundance is plotted for each sample. (E) Bar plots of the genus taxonomic levels in control, DSS+ group and DSS+CSR+ group. Relative abundance is plotted for each sample. (F) Relative abundance of genus *Odoribacter*, *Alloprevotella* and *unidentified_Lachnospiraceae* in each sample were displayed by bar plots.

**Figure S7. Flow gating strategies for each cell population.**

