## Supplementary figures and images for "Beneficial Effects of Celastrol on Immune Balance by Modulating Gut Microbiota in Dextran Sodium Sulfate-Induced Ulcerative Colitis"

### Figure S1

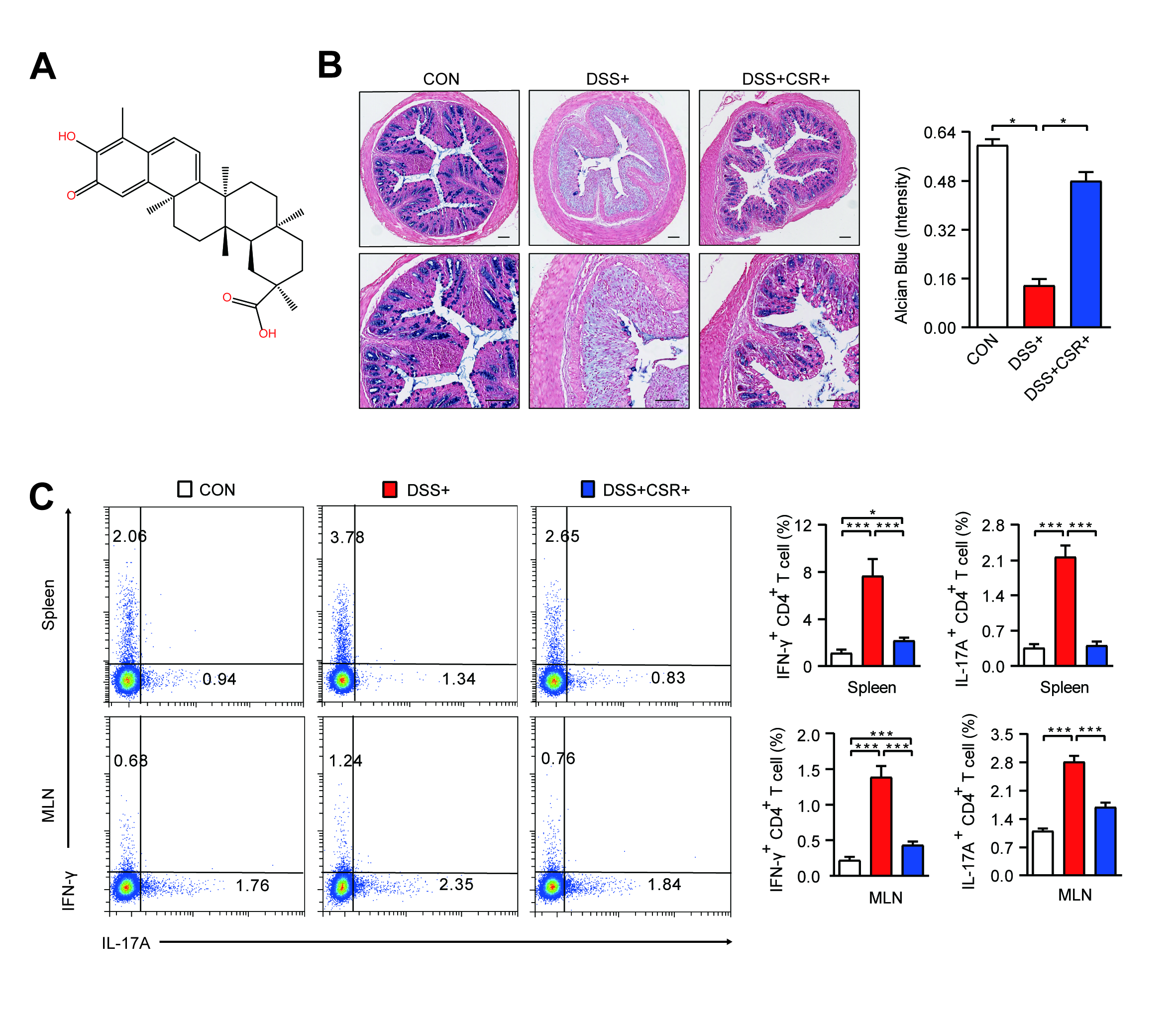

### Figure S2

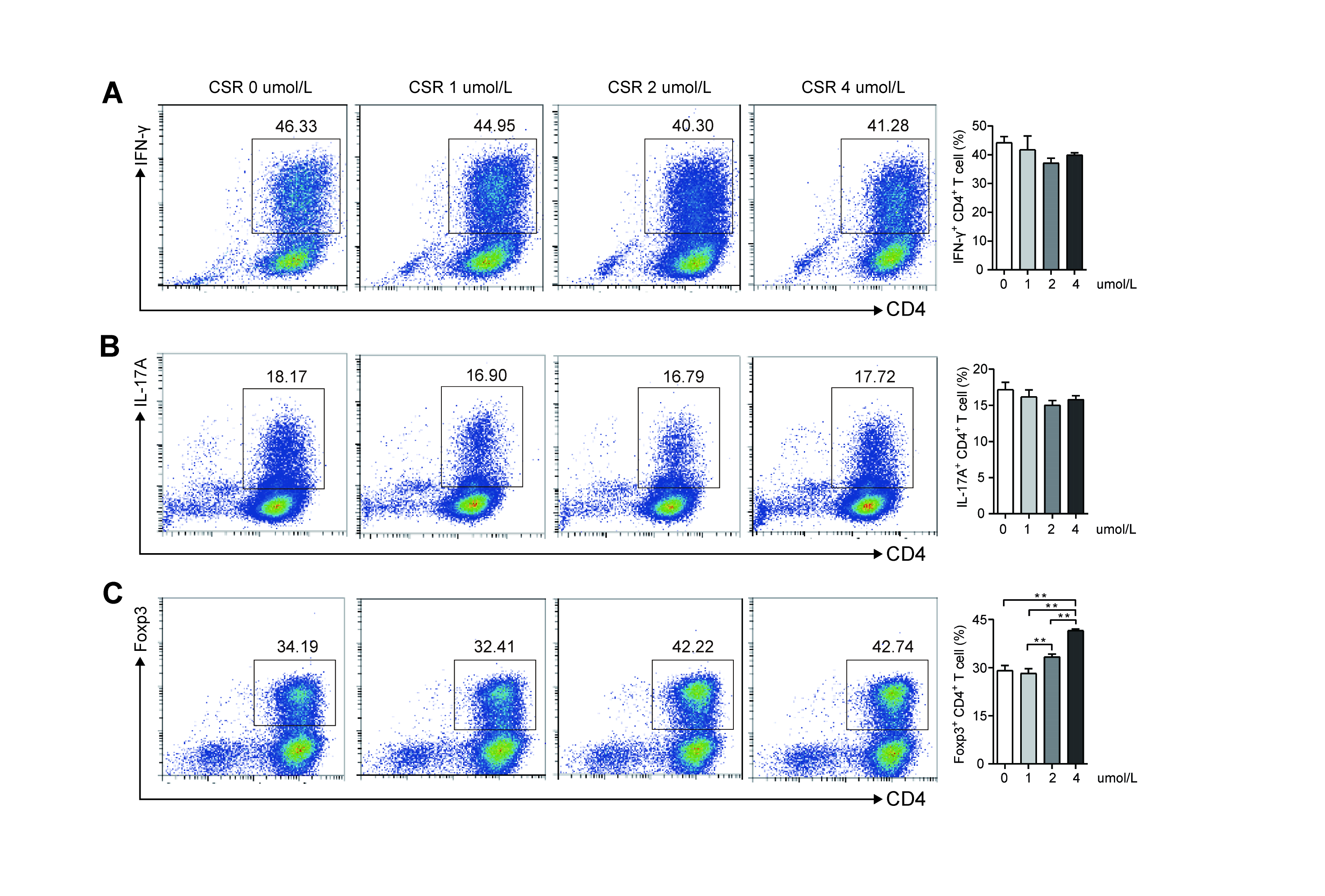

### Figure S3

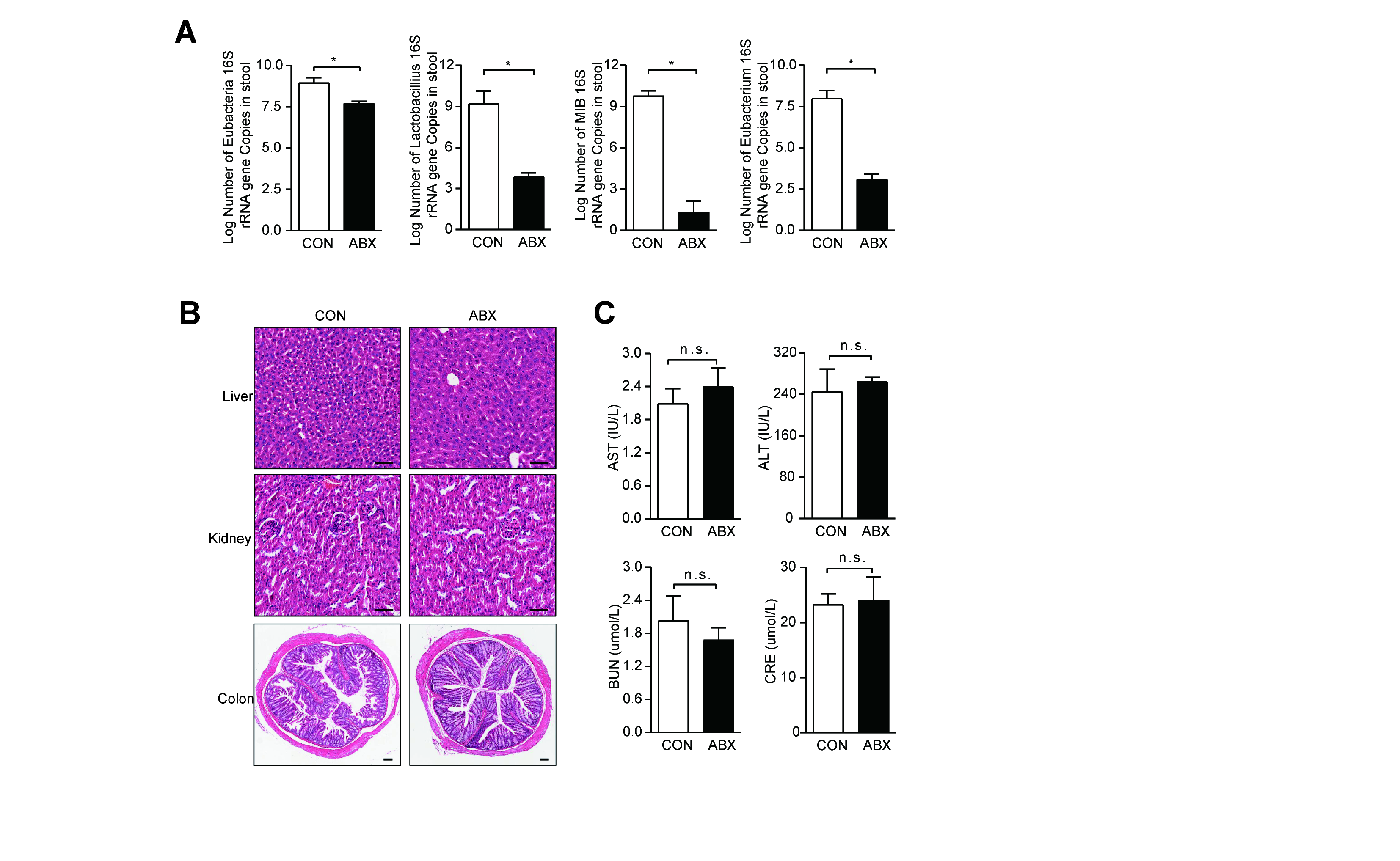

### Figure S4

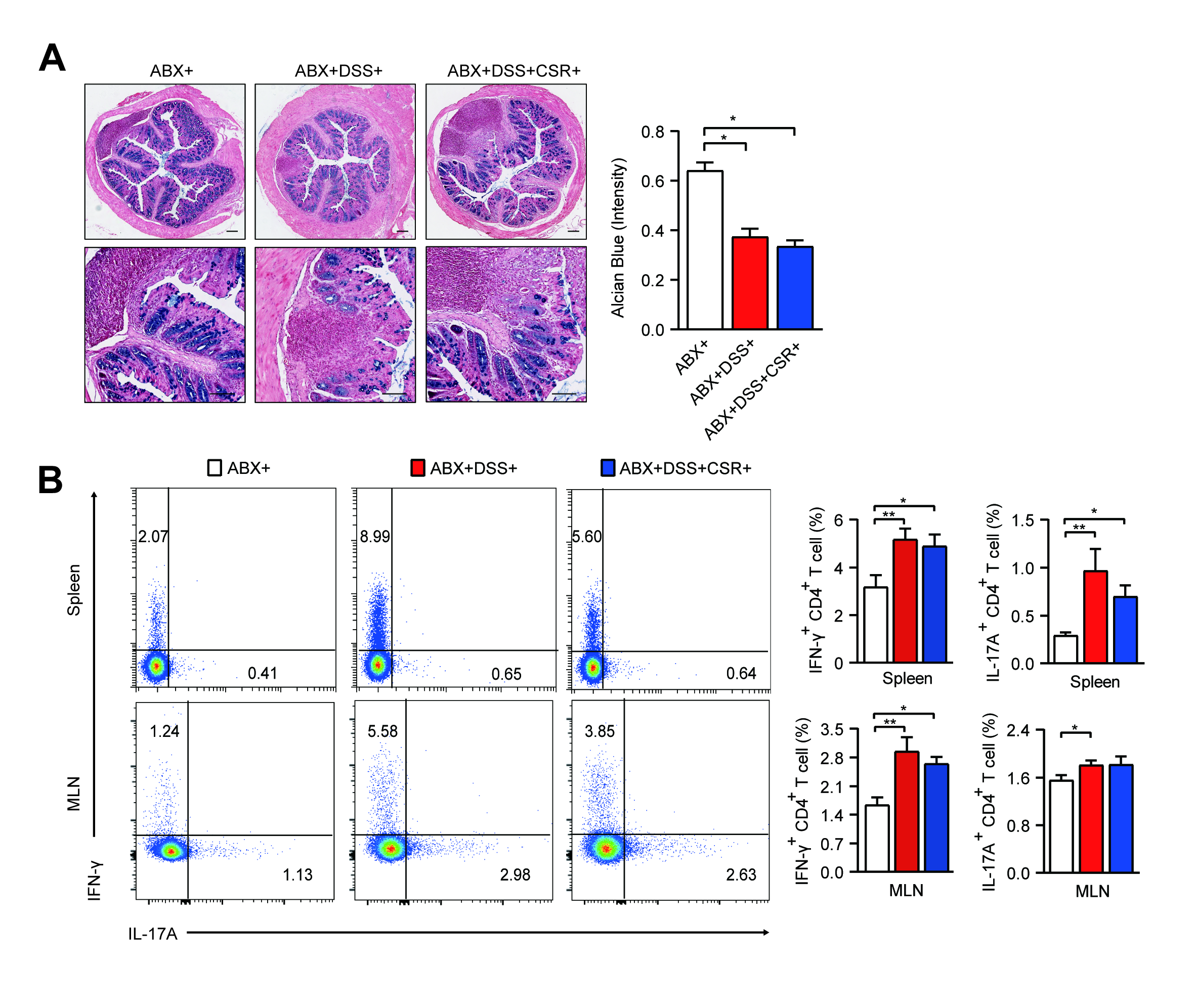

### Figure S5

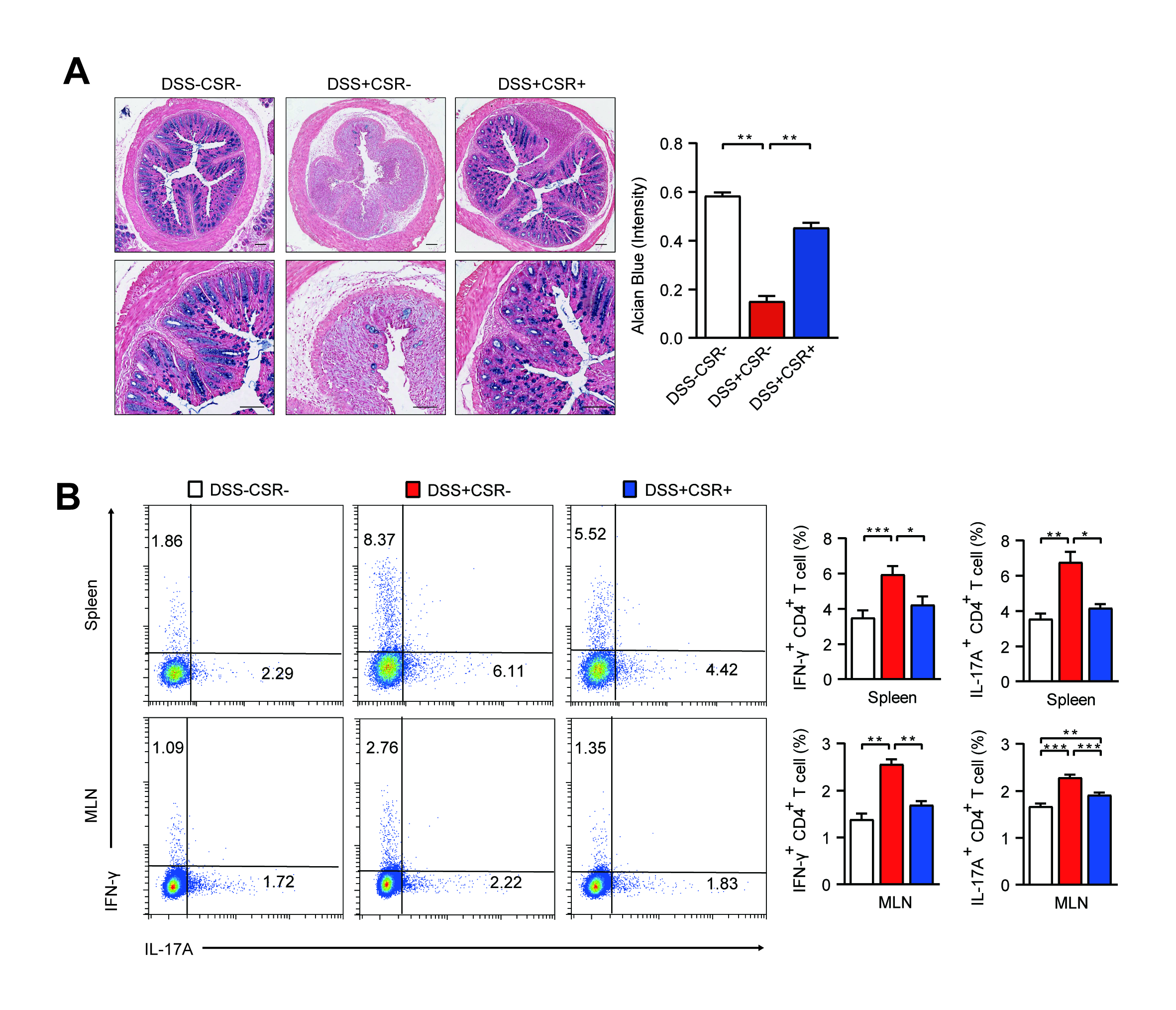

### Figure S6

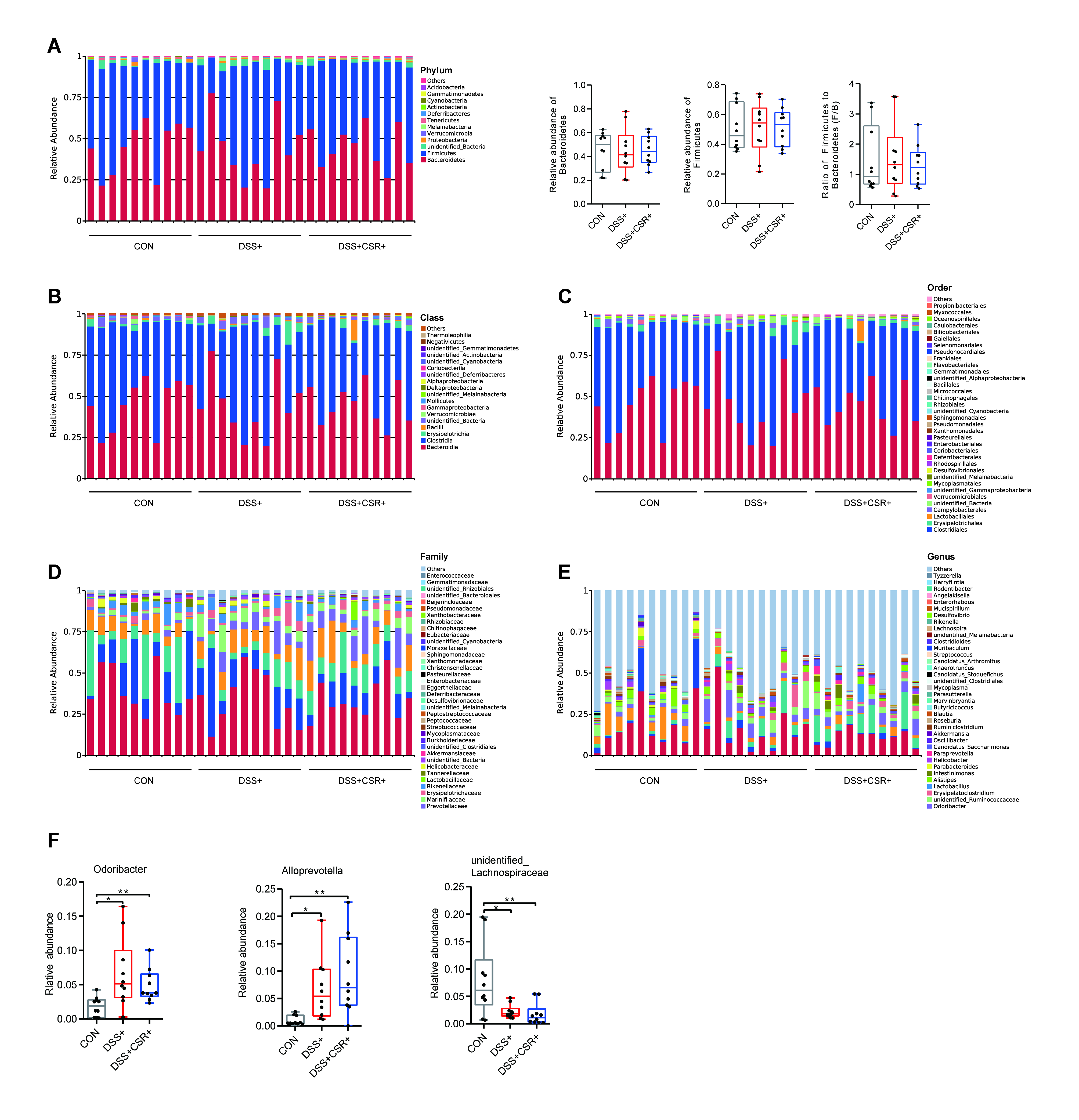

### Figure S7

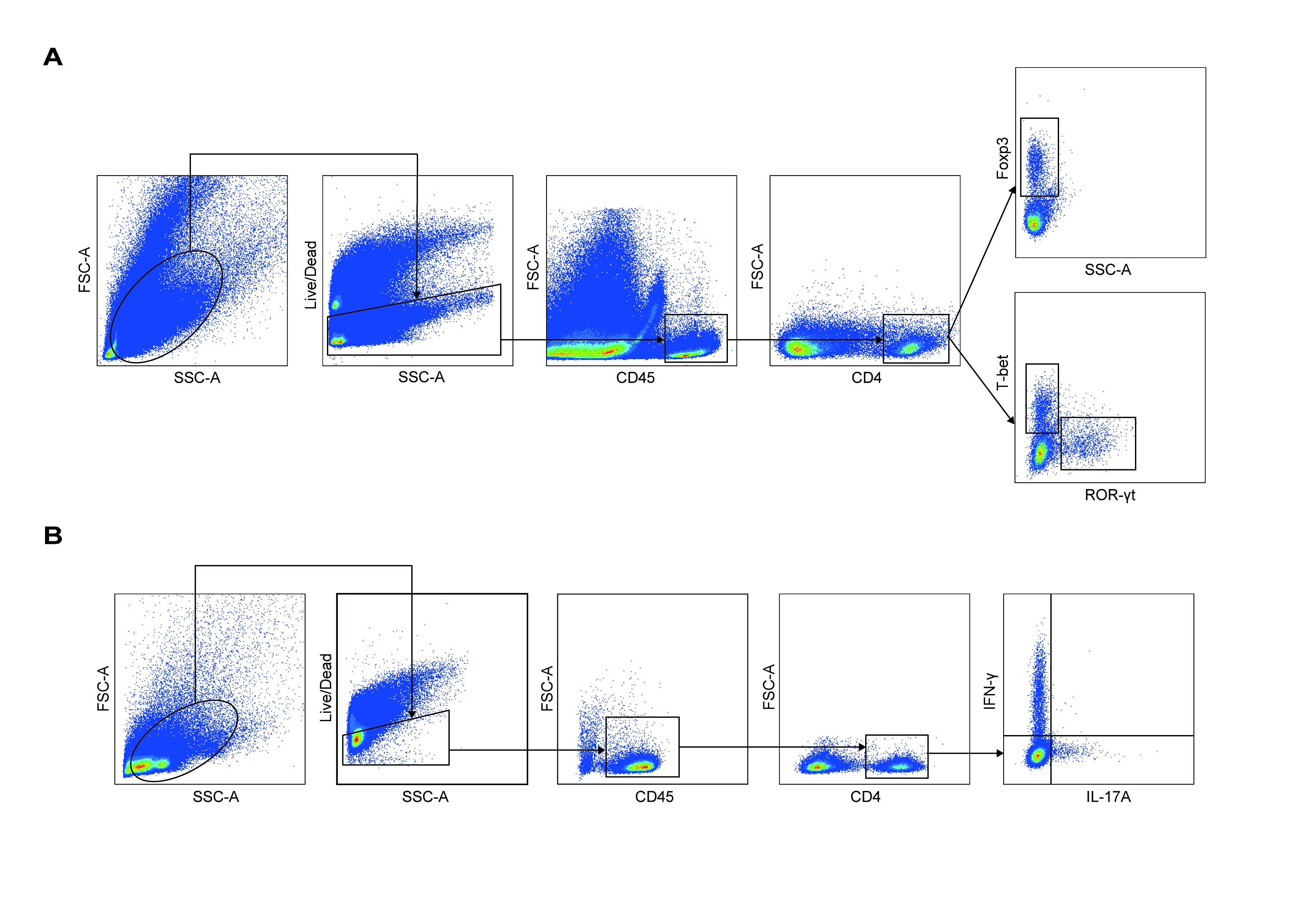
